# Shadow imaging for panoptical visualization of living brain tissue

**DOI:** 10.1101/2022.11.03.511028

**Authors:** Yulia Dembitskaya, Andrew K. J. Boyce, Agata Idziak, Atefeh Pourkhalili Langeroudi, Guillaume Le Bourdellès, Jordan Girard, Misa Arizono, Mathieu Ducros, Marie Sato-Fitoussi, Amaia Ochoa de Amezaga, Kristell Oizel, Stephane Bancelin, Luc Mercier, Thomas Pfeiffer, Roger J. Thompson, Sun Kwang Kim, Andreas Bikfalvi, U. Valentin Nägerl

## Abstract

Progress in neuroscience research hinges on technical advances in visualizing living brain tissue with high fidelity and facility. Current neuroanatomical imaging approaches either require tissue fixation, do not have cellular resolution or only give a fragmented view. Here, we show how regular light microscopy together with fluorescence labeling of the interstitial fluid in the extracellular space provide comprehensive optical access in real-time to the anatomical complexity and dynamics of living brain tissue.

---

The human brain is a structural engineering marvel where hundreds of thousands of miles worth of tightly packed axons and dendrites connect billions of neurons into a gigantic electrochemical network that generates memory, thought and action. Its main animal model, the mouse brain, is about three orders of magnitude smaller, but its anatomical structure is similarly dense and complex^1^.

Fluorescence microscopy is the method of choice for multiscale imaging of living brain tissue with high resolution. However, it typically relies on labeling sparse sets of brain cells, providing a fragmented and partial view of tissue anatomy. Electron microscopy and magnetic resonance imaging are practically label-free and unbiased approaches, yet they either offer high spatial resolution or non-invasive live imaging, but not both together. Moreover, they cannot be combined *in situ* with other powerful neuro-technologies, such as Ca^2+^ imaging, electrophysiology and optogenetics.

Getting a more detailed and broader view of brain tissue is not only important for mapping the functional connectivity of neuronal circuits^2^, it can also unearth useful information on anatomical context and tissue viability to assist in experiments and their interpretation.

Breaking this impasse, super-resolution shadow imaging (SUSHI) introduced a new paradigm to visualize the anatomy and extracellular space (ECS) of living brain tissue with nanometric spatial resolution^3^. It is based on fluorescence labeling of the interstitial fluid and super-resolution stimulated emission depletion (STED) microscopy^4, 5^, casting all membrane-enclosed cellular structures as sharply contoured ‘shadows’ against a bright background of extracellular fluorescence.

Imaging with inverted contrast is immune to bleaching because the interstitial fluid provides an inexhaustible reservoir of fresh dye molecules. It is potentially also much less invasive and harmful because the fluorophore does not need to be introduced into the cells and any phototoxic by-products do not accumulate inside of them, but can dissipate in the ECS. The most minute cellular and extracellular compartments (e.g., axon shafts, peri-synaptic astrocytic processes, spine necks, synaptic clefts) can be discerned in super-resolved shadow images^3, 6, 7^.

A recent study using the SUSHI approach and machine learning generated precise 3D reconstructions of brain tissue microstructures, even though the spatial resolution was limited to 140 nm^8^, suggesting that the neuroanatomical ground truth can be established by light microscopy when augmented by advanced computational image analysis.

Hampered by high technical demands and limited availability of super-resolution technology, SUSHI has not yet been widely adopted. Moreover, the technique has been used mostly in organotypic brain slices^6, 7^, which offer optimal imaging conditions, and not yet in acute brain slices or intact brains *in vivo*, where labeling and image contrast for shadow imaging is harder to come by.

The aim of this study was to mitigate these difficulties and make shadow imaging more versatile, outlining a straightforward (and adoptable) method for capturing more fully the complexity and dynamics of the anatomical structure of living brain tissue even without super-resolution microscopy.

To this end, we have worked out solutions for achieving innocuous, high-contrast fluorescence labeling of the interstitial fluid in acute brain slices and the intact brain *in vivo*. We show that the inverted signal can be read out by regular microscopy techniques, such as confocal, 2-photon and light sheet microscopy, providing finegrained yet expansive views of the anatomical scenery in these major experimental preparations.

We started out by imaging organotypic hippocampal brain slices using an inverted confocal microscope (**Fig. 1A**). We chose the more popular Muller slices^9^, rather than Gähwiler^10^, for which the SUSHI technique had originally been developed. Muller slices are grown on a light-scattering membrane support, but by turning it upside down and placing a metal ring on top, it is possible to get a direct and stable view from below.

**Figure 1:**
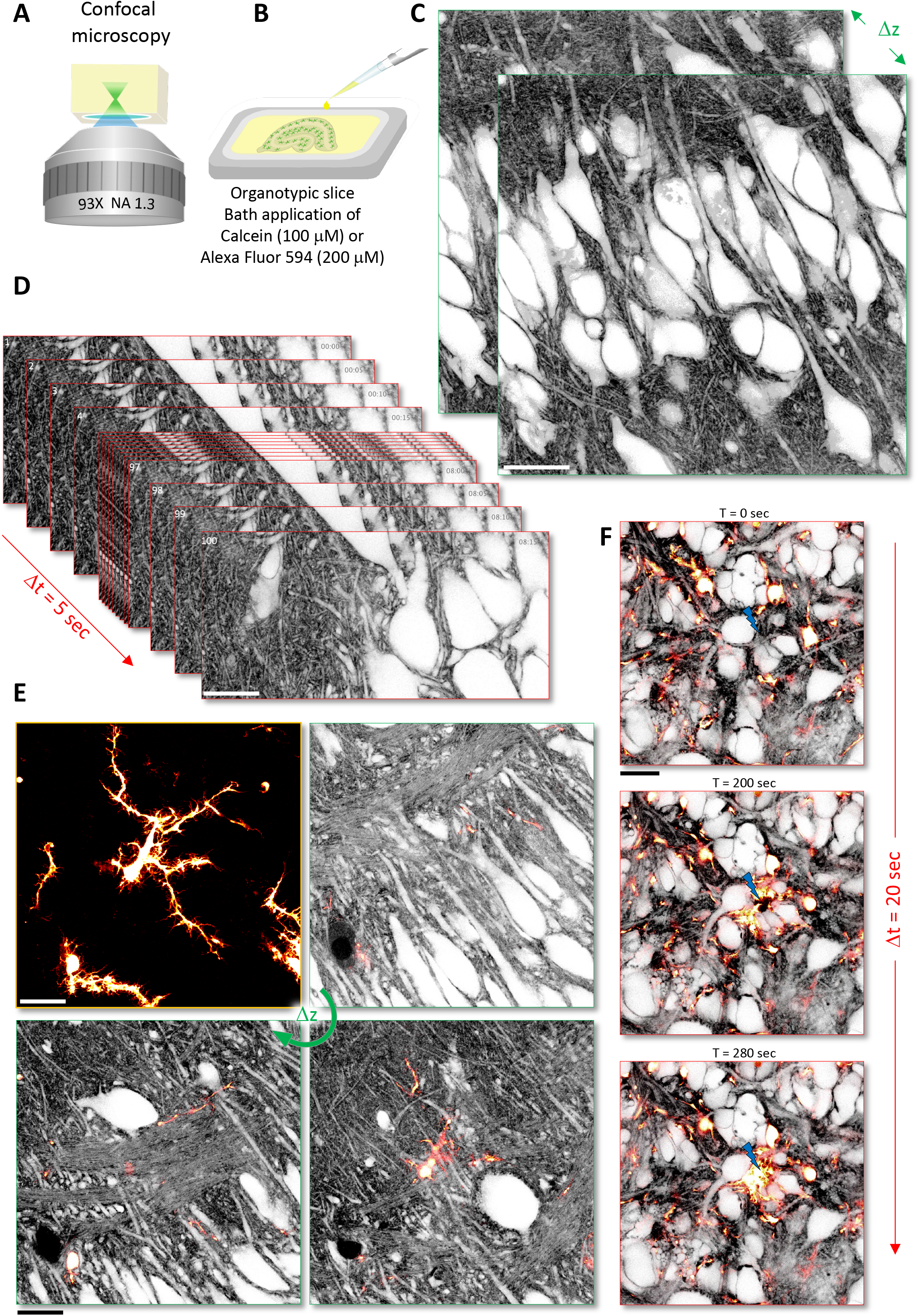
Confocal shadow imaging (COSHI) in organotypic brain slices. **(A)** Schematic of imaging technique based on a commercial inverted confocal microscope equipped with a motorized correction collar to reduce optical aberrations. **(B)** Organotypic brain slices were placed (upside down) into an imaging chamber containing the membrane-impermeant organic dye diluted in ACSF (100 μM of Calcein, or 200 μM of Alexa Fluor 594). **(C)** Z-stack of confocal shadow images (Calcein, 100 μM) of CA1 area of hippocampus in an organotypic brain slice (see also **SI Video 1A, B** and **C**). Scale bar, 20 μm. **(D)** Time series of confocal shadow images (Calcein, 100 μM) of CA1 area of hippocampus in an organotypic brain slice, 1 frame every 5 seconds, 100 frames in total (see also **SI Video 2**). Scale bar, 20 μm. **(E)** Z-stack of confocal shadow images (AlexaFluor 594, 200 μM) of CA1 area of hippocampus in an organotypic brain slice from a transgenic mouse line (CX3CR1-GFP), where all microglial cells are fluorescently labeled with GFP (see also **SI Video 3**). First panel shows maximum-intensity projection of only GFP-labeled microglia. Other panels show overlay of GFP and shadow image for three different optical sections. Scale bars, 20 μm. **(F)** Time series of confocal shadow images (AlexaFluor 594, 200 μM) showing labeled microglia cells (CX3CR1-EGFP) that react to a laser lesion (blue bolt), rapidly extending their fine processes towards the site of the lesion (see also **SI Video 4**). Scale bar, 20 μm.

We used a 93X glycerol-immersion objective (NA 1.3) equipped with a motorized correction collar to reduce spherical aberrations caused by the mismatch in refractive index between medium (*n* ∼ 1.46) and brain slice (*n* ∼ 1.37). The spatial resolution of the microscope was around 212 nm in x-y and 550 nm in z (data not shown).

To generate morphological contrast, we added a small but membrane-impermeant organic fluorescent dye (Calcein, 100 μM, **Fig. 1B**) to the ACSF bath solution in which the brain slices were submerged, as described before^3^. After optimizing the correction collar, we could acquire images of the tissue with high contrast and resolution (**Fig. 1C**), offering a macroscopic perspective of the anatomical layout replete with crucial structural details, such as dendrites and axons.

Despite the high density of the fluorescent label, it was possible to acquire confocal shadow images (COSHI) and z stacks at least 50 μm below slice surface (**SI Video 1A, B, C**), before image quality decreased due to out-of-focus blur, aberrations and light scattering.

Given the diffusional replenishment of dye molecules, it was possible to acquire a high number (>100) of time-lapse frames (**Fig. 1D**; **SI Video 2**) with little or no drop in signal-to-noise ratio (SNR) or signs of phototoxicity (data not shown).

Next, we checked the utility of COSHI for studying microglia and their morphological dynamics in living brain tissue. Microglia are tissue-resident macrophages that are critical for the immune defense of the brain. They have highly branched and motile processes, which touch and seem to influence dendritic spines during brain development and neuroplasticity^11, 12^. However, it remains unclear, which other cellular structures they come into contact with in the surrounding neuropil.

To this end, we used organotypic brain slices from transgenic mice (CX3CR1-EGFP) where microglia are highlighted by GFP and labeled the ECS with a red dye (Alexa Fluor 594, 200 μM). Indeed, COSHI made it possible to reveal the complex arborization of microglial processes amidst their fully visible anatomical context (**Fig. 1E** and **SI Video 3**). Moreover, after inflicting a local laser lesion, which triggers a rapid and orchestrated immune reaction, it was possible to see how microglial processes navigate through the dense anatomical landscape towards the lesion site (**Fig. 1F, SI Video 4**). Thus, the shadow imaging approach paired with positive labeling may facilitate the study of cell growth and motility in various tissue micro-environments.

As a point-scanning technique, it typically takes several seconds to acquire a single confocal image, which is often too slow for imaging Ca^2+^ transients and other dynamic biochemical activities. In contrast, light-sheet microscopy reconciles high spatial with high temporal resolution because fluorescence excitation and detection are orthogonal to each other, enabling fast widefield imaging without out-of-focus blur (**Fig. 2A**)^13^.

**Figure 2:**
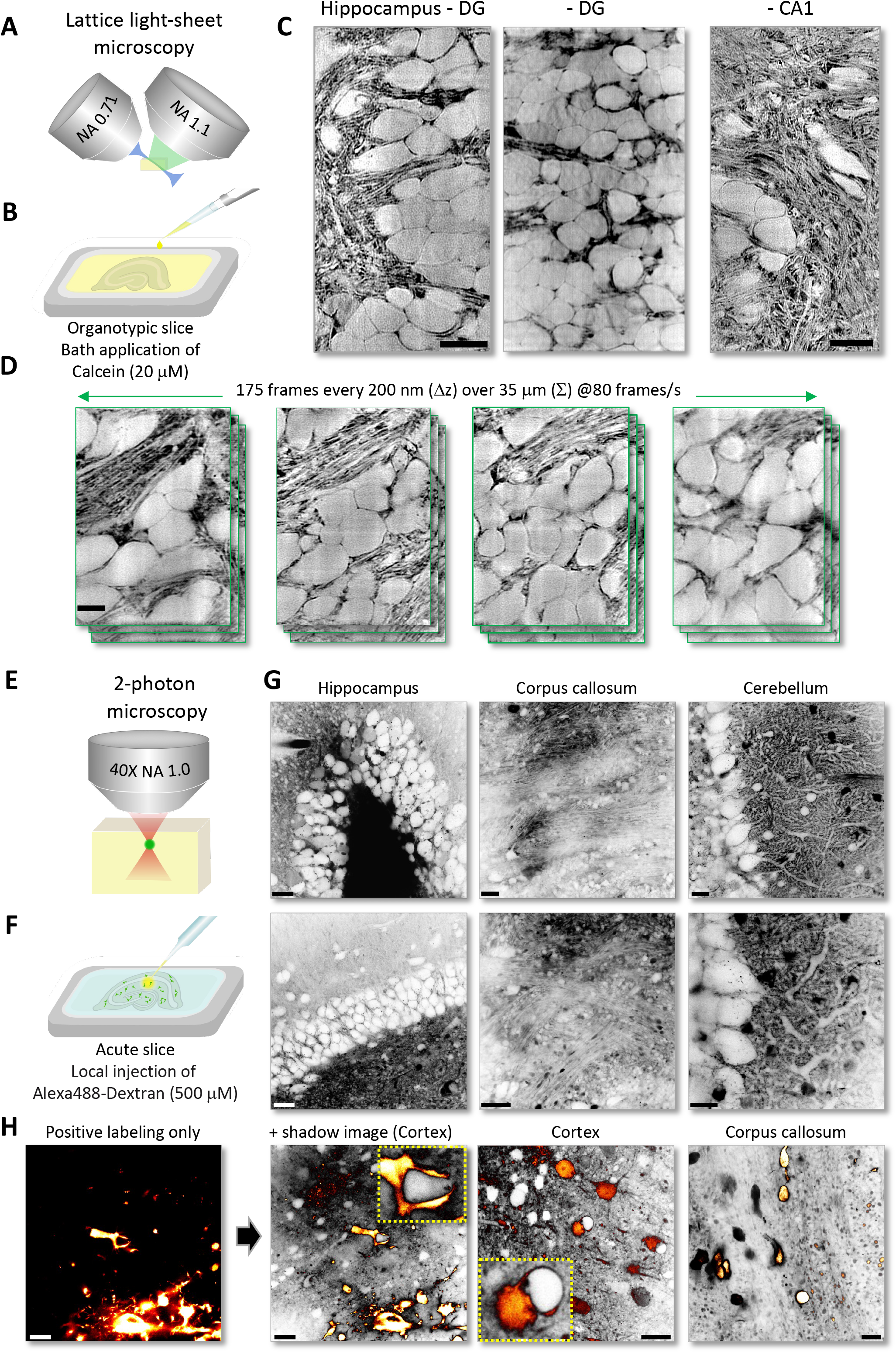
Shadow imaging with light-sheet (LISHI) and 2-photon microscopy (TUSHI) in brain slices. **(A)** Schematic of imaging technique based on a custom-built lattice light-sheet microscope. **(B)** Organotypic brain slices were placed (right side up) into an imaging chamber containing the membrane-impermeant organic dye diluted in ACSF (20 μM of Calcein). **(C)** Shadow images acquired by light-sheet microscopy in different areas of the hippocampus in organotypic brain slices (see also **SI Video 5** and **6**). Scale bar, 10 μm. **(D)** High-speed z-stack image acquisition in organotypic brain slice, 175 frames acquired at 80 Hz and a Δz step size of 200 nm. Scale bar, 5 μm. **(E)** Schematic of imaging technique based on a commercial 2-photon microscope. **(F)** Acute brain slices were placed into the imaging chamber and a micro-manipulated patch pipette was used to inject the extracellular dye well below the surface of the slice (500 μM of Alexa488-Dextran). **(G)** Representative 2-photon shadow images of acute brain slices from different regions including hippocampus, corpus callosum and cerebellum, revealing their characteristic anatomical structure. Scale bars, 20 μm. **(H)** Shadow imaging of fluorescently labeled tumor cells in acute brain slices that were implanted into the brains of mice four weeks before slicing, revealing their anatomical relationship with the cellular environment, examples from cortex and corpus callosum. Scale bars, 20 μm. Please note instances of intimate ‘embrace’ of unlabeled structures/cells by tumor cells.

To explore if shadow imaging is compatible with light-sheet microscopy, we used a custom-built microscope to image organotypic brain slices, where a thin and homogenous light sheet was created by a lattice excitation pattern and an oscillating mirror, as described previously^14, 15^. The spatial resolution of the system was around 273 nm in x-y and 524 nm in z (data not shown).

Because of the efficient signal detection scheme in light-sheet microscopy, it was possible to use much lower dye concentrations (Calcein, 20 μM, **Fig. 2B**). We could acquire high-contrast shadow images of the tissue, revealing fine details of its cellular microarchitecture (**Fig. 2C**), almost as well as COSHI but with >100X higher imaging speeds (**Fig. 2D, SI Video 5** and **6**). Thus, light-sheet shadow imaging (LISHI) of cellular structures can in principle be performed alongside high-speed imaging of biochemical dynamics such as synaptic release of glutamate and its spread in the ECS.

2-photon microscopy is the main technique used for imaging acute brain slices or *in vivo*, offering superior SNR and optical sectioning deep inside light scattering tissue (**Fig. 2E**)^16^. However, shadow imaging in these popular experimental preparations poses unique challenges for fluorescence labeling and image contrast. Inevitably, there are many dead and cut open cells on the surface of an acute slice, which will take up the fluorescent dye if it is bath-applied, diminishing image contrast between cellular compartments and ECS.

To circumvent this problem, we spritz-injected the dye inside of the brain slices (>50 μm below surface), where cells are mostly intact, via a pressurized patch pipette (**Fig. 2F**). Using a commercial 2-photon microscope with a 40X water-immersion objective (NA 1.0, spatial resolution x-y = 350 nm and z = 1.5 μm; data not shown), it was also possible to obtain shadow images with high contrast over large fields of view, temporarily lighting up the anatomical layout of the slices (**Fig. 2G**).

To increase and prolong fluorescence contrast, we used a dextran-conjugated dye (Alexa Fluor 488-Dextran, 500 μM), which slowed the dispersion of the dye in the ECS, making it possible to inject a relatively small volume (<1 μL) at low pressure with minimal tissue disturbance.

To demonstrate the ability of 2-photon shadow imaging (TUSHI) to reveal the anatomical context of a specific set of fluorescent cells, we spritzed the dye into acute brain slices prepared from mice implanted with YFP-labeled GBM tumor cells, which is a mouse model of glioblastoma of the mesenchymal subtype^17, 18^. With two fluorescence detection channels (for YFP and Alexa Fluor 488), it was possible to image the proliferating tumor cells and appreciate their spatial integration in the tissue (**Fig. 2H**). Of note, this is the first study where the shadow technique is applied to tumor tissue.

Taken together, the ‘spritz-shadow imaging’ technique, which is a variant of ‘shadow-patching’ for targeted electrophysiological recordings *in vivo*^*19*^, can on the fly enrich slice physiology studies with visual information on anatomical context.

Finally, we set out to extend the shadow imaging concept to the mouse brain *in vivo* to pave the way towards longitudinal neuroanatomical studies in mouse models of neuroplasticity and brain diseases.

We used a home-built 2-photon microscope with a 60X silicone oil objective (NA 1.3, WD 0.3 mm) that was equipped with a programmable spatial light modulator (SLM) to help reduce optical aberrations when imaging deep inside brain tissue (**Fig. 3A**). The spatial resolution of the microscope was around 320 nm in x-y and 925 nm in z (data not shown).

**Figure 3:**
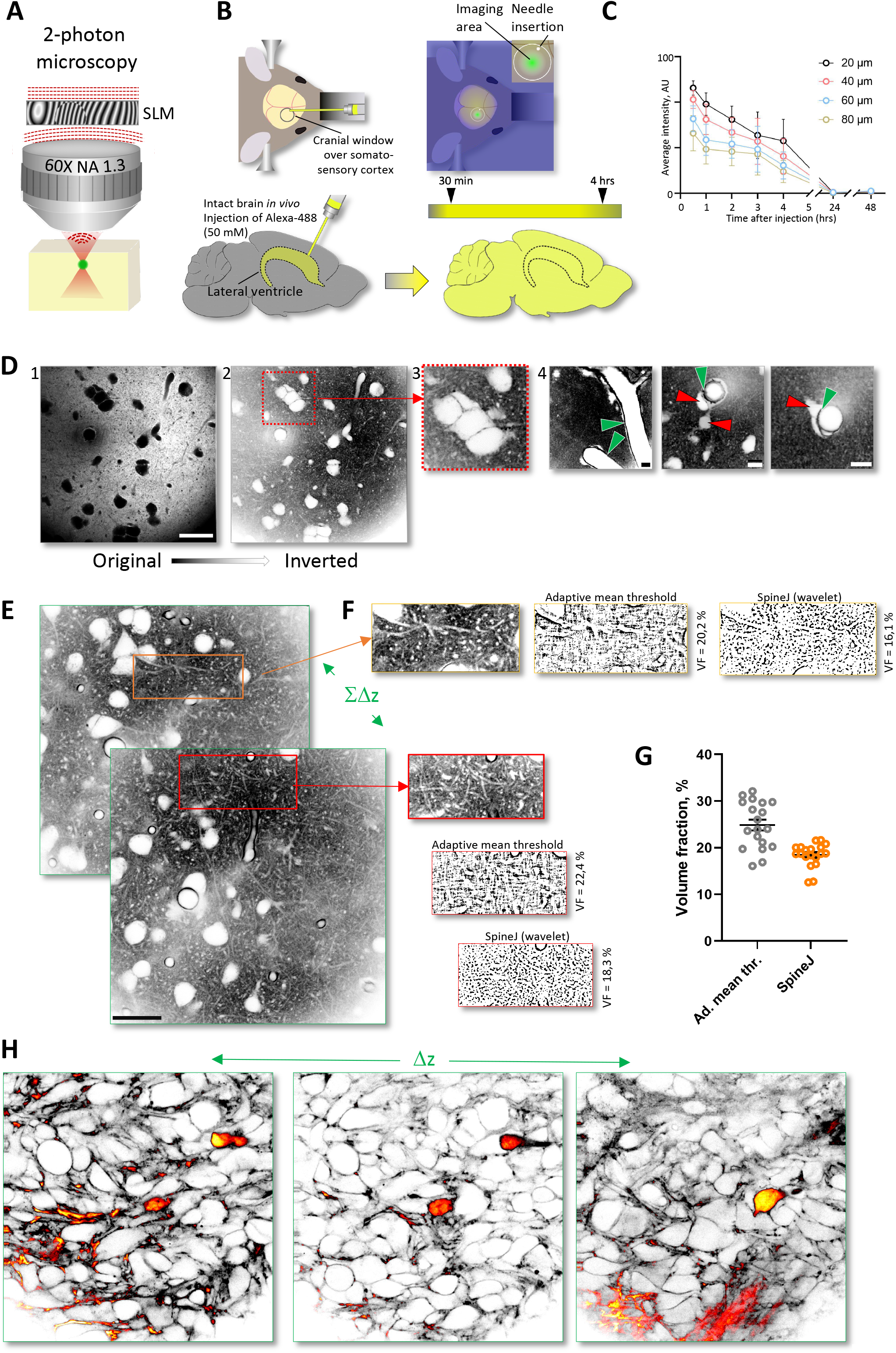
Shadow imaging with 2-photon microscopy in the mouse brain *in vivo*. **(A)** Schematic of imaging technique based on a custom-built 2-photon microscope equipped with a spatial light modulator (SLM) and a high-NA objective with a correction collar to reduce optical aberrations. **(B)** Schematic of the *in vivo* labeling strategy of the ECS, where a small volume of highly concentrated dye solution (8 μL of 50 mM of Alexa Fluor 488) was stereotaxically injected slowly (over 10 min) into the ipsilateral lateral ventricle. **(C)** Time course of average fluorescence labeling intensity of the ECS after intraventricular dye injection, measured at different depths below the cortical surface. The fluorescence labeling appeared within 30 minutes and stayed elevated at workable levels for at least 4 hours. **(D)** Representative *in vivo* 2-photon shadow images acquired in the somatosensory cortex of an anesthetized mouse. Panel 1 (left): raw image, revealing cell bodies as large black ‘shadows’; Scale bar, 25 μm; Panel 2: same image after inversion of the look-up-table. Panel 3: zoomed-in part showing cluster of cell bodies; Panel 4: Examples of shadow images delineating perivascular spaces (green arrows) running along blood vessels and putative pericytes and/or astrocytic endfeet (red arrows) wrapping around blood vessels. Scale bars, 10 μm. **(E)** Z-stack of 2-photon shadow images acquired over >100 μm in depth (see also **SI Video 7**). Scale bar, 25 μm. **(F)** Regions of interest in neuropil analyzed for ECS volume fraction, using two different algorithms for binarizing the images (based on ‘adaptive mean thresholding’ or ‘wavelet filtering’ using SpineJ). **(G)** Group data of ECS volume fraction of cortical neuropil, indicating mean values of around 20-25%, consistent with biophysical measurements in the literature. **(H)** *In vivo* 2-photon shadow images in motor cortex of mice implanted with YFP-labeled cells from a human GBM tumor cell line. The Alexa Fluor 488 (ECS) and YFP (tumor cells) fluorescence signals were spectrally detected. The images shown here were merged and pseudo-colored based on signal intensity.

In addition to the usual challenges of *in vivo* imaging, such as brain motion and limited optical access, shadow imaging *in vivo* requires generating fluorescence contrast inside a large and sealed-off compartment. The labeling of the interstitial fluid should be strong, long-lasting and minimally invasive. Instead of injecting the dye directly into the tissue, which proves to be too disruptive and unreliable, we did stereotaxic injections into the ipsilateral ventricle controlled by a high-precision syringe pump, delivering the dye (∼8 μL of 50 mM of Alexa Fluor 488) over an extended period of time (∼10 min) (**Fig. 3B**). This amount theoretically corresponds to a dye concentration of a few millimolar in the interstitial fluid, if averaged over the entire mouse brain ECS. However, because of clearance, the effective concentration likely was much lower.

In this way, it was possible to achieve reproducible and strong brain-wide fluorescence ECS labeling that peaked shortly after the injection (<1 hour) but persisted at workable levels for at least four hours (**Fig. 3C**). The procedure was optimized for the animals to recover and live on normally.

Following standard protocols for craniotomy and window implantation, leaving the dura mater intact, we could create stable and clear optical access to the mouse cortex. An anesthesia protocol based on intraperitoneal injections of ketamine and xylazine, and careful physical positioning of the animal under the microscope using ear-bars for head fixation prevented almost all motion artifacts from inspiratory muscle contractions, enabling image acquisitions with minimal motion blur.

Applying these technical measures, we could acquire high-contrast 2-photon shadow images of the superficial layers of somatosensory cortex (**Fig. 3D**). The resolution and contrast of the images were lower for *in vivo* than for organotypic brain tissue, which can be explained by the use of longer wavelength light 2-photon excitation (λ = 920 nm *versus* 488 nm), residual brain motion and presumably lower dye concentrations *in vivo* due to clearance from the interstitial fluid. Nevertheless, the TUSHI approach can reveal the outlines of neuronal cell bodies and blood vessels, where finer details like astrocytic endfeet, pericytes and perivascular spaces become readily visible.

Moreover, by taking z stacks of up to 100 μm in depth, it was possible to survey sizable volumes of brain tissue (**Fig. 3E**; **SI Video 7**). Because the dye is diffusible in the ECS, it was possible to acquire a high number of time-lapse images unaffected by bleaching (data not shown).

The volume fraction (VF) indicates the relative amount of ECS in the brain, estimated to be about a fifth of the entire brain. It is an important structural parameter of the tissue, which can be measured precisely by biophysical techniques but typically only over large tissue volumes^20^. By contrast, the VF can be determined directly anywhere in a field of view at micron scale by distinguishing ECS from cellular structures in shadow images simply by image binarization (**Fig. 3F**). We estimated the VF of cortical neuropil to be around 20-25% based on two different algorithms for image binarization (**Fig. 3G**). These values are much higher than what EM images of chemically fixed brain tissue suggest, but in line with the biophysical literature.

To confirm that TUSHI *in vivo* can be combined with imaging positively labeled cells, we again turned to the mouse model of glioblastoma^17, 18^. The approach showed YFP-labeled tumor cells infiltrating a cluster of neuronal cell bodies in motor cortex four weeks after their implantation into the mouse brain (**Fig. 3H**).

The technique may be valuable for studying various primary or secondary brain tumors in mouse models^21^, how tumor cells interact with tissue microenvironment and impact brain structures^22^. It also is uniquely suited for *ex vivo* tissue samples where positive labeling is not an option, such as fresh human pathology specimen from the clinics, as a diagnostic tool or for monitoring therapy. In principle, TUSHI could be combined with any other form of positive cell labeling, ie. circuit tracing, cell type reporters, or fluorescent biosensors to explore brain structure and function in view of the complete cellular context in living animals.

Taken together, our study presents a strong case for shadow imaging of living brain tissue using common diffraction-limited microscopy techniques. It can help reveal the anatomical context of labeled cellular players in a variety of patho-physiological settings, in addition to providing ECS structural information. In view of ongoing advances in ECS labeling, super-resolution microscopy and computational tools, shadow imaging is bound to become an indispensable tool for live-tissue microscopy.

## Acknowledgements

The study was supported by grants from the European Research Council (ERC-SyG ENSEMBLE #951294), Human Frontiers Science Program (#RGP0036/2020), ERA-NET NEURON (ANR-17-NEU3-0005) and Agence Nationale de la Recherche (ANR-17-CE37-0011) to UVN. AKJB was funded by an Alberta Innovates Postdoctoral Fellowship (2+1), a EuroBio Imaging User Access Fund Award, and a Rebecca Hotchkiss International Scholar Exchange Award. AI was funded by a fellowship from Bordeaux Neurocampus Graduate Program. SKK was supported by a National Research Foundation of Korea grant (NRF-2019R1A2C2086052). We thank the animal facilities at IINS and in Talence for their professional support.

## Author contributions

AKJB, AI, AP and YD collected all imaging data. AKJB, AI, AP, MA, MD, UVN, YD analyzed the data and prepared figures. AKJB, AI, AOA, AP, GL, JG, LM, KO, MA, MD, MSF, SB, SKK, TP, UVN and YD provided technical input for sample preparation, labeling and microscopy. KO and AB provided reagents. AB and RJT provided postdoc supervision. UVN conceived and supervised the study and wrote the paper with input from co-authors.

## Competing financial interest

The authors declare no competing financial interests.

## Supplemental Videos

**SI Video 1A, B, C: COSHI z stacks in organotypic brain slices**

**SI Video 2: COSHI time series**

**SI Video 3: COSHI z stack showing GFP-labeled microglia cell**

**SI Video 4: COSHI time series showing microglial response to laser lesion**

**SI Video 5: LISHI z stack**

**SI Video 6: LISHI time series of z stack**

**SI Video 7: TUSHI z-stack *in vivo***

**SI Video 8: TUSHI z stack showing YFP-labeled tumor cells in motor cortex *in vivo***

## Online Methods

### Animals / brain tissue samples

#### Regulatory issues

Animal handling and experimental procedures were in accordance with the European Union and CNRS institutional guidelines for the care and use of laboratory animals (Council directive 2010/63/EU) and approved by the Institutional Animal Care and Use Committee at the University of Bordeaux, France (DAP 2019031909389750_v7).

#### Mouse lines

C57Bl/6J wild-type, CX3CR1-EGFP and immunodeficient Rag 2 Gamma C-/-mice were used in this study. Mice were housed under a 12 h light/12 h dark cycle at 20-22 °C with *ad libitum* access to food and water in the animal facility of the Interdisciplinary Institute for Neuroscience and of Animalerie Mutualisée de Talence (University of Bordeaux/CNRS) and monitored daily by trained staff. All animals used were free of any disease or infection at the time of experiments. Pregnant females and females with litters were kept in cages with one male. We did not distinguish between males and females among the perinatal pups used for organotypic cultures, as potential anatomical and/or physiological differences between the two sexes were considered irrelevant in the context of this study.

#### Organotypic brain slices

Organotypic slice cultures were prepared according to the Muller method^9^. Briefly, hippocampal slices were obtained from postnatal day 5-7-old C57BI/6J mouse pups. The animals were quickly decapitated, and the brains placed on cold sterile dissection medium (all in mM; 0.5 CaCl_2_, 2.5 KCl, 2 MgCl_2_, 0.66 KH_2_PO_4_, 0.85 Na_2_HPO_4_-12H_2_O, 0.28 MgSO_4_-7H2O, 50 NaCl, 2.7 NaHCO_3_, 25 glucose, 175 sucrose, 2 HEPES; all from Sigma). The hippocampi were dissected and sliced on a McIlwain tissue chopper to generate coronal brain slices of 350 μm thickness. After 20 minutes of incubation at 4°C, the slices were transferred onto sterilized hydrophilic polytetrafluoroethylene (PTFE) membrane (FHLC04700; Merck Millipore) pieces, which were placed on top of cell culture inserts (Millipore, 0.4 mm; Muller method); the inserts were held in a 6-well plate filled with medium (50% Basal medium eagle (BME), 25% Hank’s Balanced Salt Solution (HBSS), 25% Horse Serum, 11.2 mmol/L glucose and 20 mM glutamine; all from GIBCO) and cultured up to 14 days at 35°C/5% CO_2_. For CX3CR1-EGFP slices, culture medium was replaced with microglial-supportive growth medium (50% Basal medium eagle (BME), 36.5% Hank’s Balanced Salt Solution (HBSS), 10% Horse Serum, 2% B27 plus supplement, 11.2 mmol/L glucose and 20 mM glutamine; all from GIBCO Culture) at DIV3-4. Culture medium was replaced every two days.

Organotypic slices were imaged in HEPES-buffered ACSF maintained at 34 ºC or 37ºC (all in mM: 119 NaCl, 2.5 KCl, 1.3 MgSO_4_, 1 NaH_2_PO_4_-2H_2_O, 20 D-glucose, 1.5 CaCl_2_, 10 HEPES; pH 7.4, 300 mOsm; all salts from Sigma).

#### Acute brain slices

Acute mouse brain slices were prepared (coronal and sagittal). Mice were anesthetized with isoflurane or ketamine/xylazine (100/10 mg/kg) and perfused intracardially with ice-cold (for cortical and hippocampal slices) or warm (34°C, for cerebellar slices) NMDG-based solution and then decapitated. NMDG-based solution contained (in mM): 92 NMDG, 2.5 KCl, 1.25 NaH_2_PO_4_, 30 NaHCO_3_, 20 HEPES, 25 glucose, 2 Thiourea, 5 Na-ascorbate, 3 Na-pyruvate, 0.5 CaCl_2_·2H2O, and 10 MgSO_4_·7H_2_O. Extracted brains were placed in either ice-cold or warm NMDG-based solution to prepare 350 μm-thick slices using a vibrating microtome (Leica VT1200S, Leica Microsystems). Slices were transferred to NMDG-based solution at 34-36ºC for 15-30 min for initial recovery and then stored in a solution containing (in mM): 125 NaCl, 2.5 KCl, 10 glucose, 1.25 NaH_2_ PO_4_, 2 sodium pyruvate, 3 myo-inositol, 0.5 sodium ascorbate, 26 NaHCO_3_, 2 CaCl_2_, 1 MgCl_2_ for 30 minutes at 34-37ºC and at room temperature afterwards.

For imaging, the slices were placed into a microscope chamber maintained at 32 ºC where they were continuously perfused at a rate of 2 ml/min with an ACSF solution containing (in mM): 125 NaCl, 2.5 KCl, 11 glucose, 25 NaHCO_3_, 1.25 NaH_2_PO4, 2 CaCl_2_, 1 MgCl_2_. The ACSF had an osmolarity of 295-300 mOsm, a pH of 7.4 and was continuously carbogenated (95% O_2_ and 5% CO_2_).

#### Brains in vivo

For *in vivo* imaging, we used 3-5 months old C57B6/J or 3-4 months-old Rag 2 Gamma C-/-mice of both sexes. Animals were injected with buprenorphine (0.1 mg/kg) prior to the surgery for pain relief. Surgery was done under isoflurane anesthesia. A round craniotomy (∼4 mm in diameter), which left the dura mater intact, was made above somatosensory (C57B6/J mice) or motor M1 (Rag 2 Gamma C-/-mice) cortex. The dye (Alexa Fluor™ 488, carboxylic acid, Invitrogen, 8-10 μl with a concentration of 20 or 50 mM) was injected into the lateral ventricle on the side of the craniotomy at coordinates: M/L-1.1, A/P – 0.52, D/V – 2.3 on the side of craniotomy at a rate of 1μl/min using a motorized syringe pump. The craniotomy was covered with a glass coverslip (#1 thickness, diameter - 4 mm) and sealed with glue and dental cement (Superbond C&B). After surgery, mice were anesthetized with ketamine/xylazine (100/10 mg/kg) and placed on a heated blanket under the objective of a 2-photon microscope.

### Implantation of YFP-labeled glioblastoma cells

Orthotopic injections of 104 human YFP-positive GBM cells into 6-8 weeks old immuno-deficient laboratory mice (Rag 2 Gamma C-/-) were performed using a stereotactic frame at 2□mm on the left of the medial suture and 3□mm deep in the striatum.

The P3 primary GBM cell line was derived from a tumor patient with high grade glioma^17^. After resection of a small piece of the tumor, it was mechanically dissociated (Miltenyi) and grown in NBM media (ThermoFischer) supplemented with 2□mM l-glutamine, B27 supplement (ThermoFischer), 2□μg/ml heparin (Sigma Aldrich), 20□ng/ml EGF (Peprotech) and 25□ng/ml bFGF (Peprotech), 100□U/ml penicillin (Sigma Aldrich), and 100□μg/ml streptomycin (Sigma Aldrich). The P3 cell line was transfected with a lentiviral plasmid FUGW-EYFP to express YFP in the cytosol (Addgene).

### Confocal microscopy

Muller organotypic slices were imaged using a commercial confocal microscope (Leica-SP8-STED Falcon) with a 93X glycerol immersion objective (NA 1.3) equipped with a motorized correction collar (Leica Microsystems). The membrane support holding the slices was inverted and placed on a circular coverslip (18 mm diameter, # 1.5) in a Ludin imaging chamber (Life Imaging Services), so that the slice directly faced the coverslip. A brass ring was placed on top of the membrane to prevent it from drifting. For wildtype slices, Calcein (Dojindo Laboratories) was diluted to a final concentration of 100 μM in the ACSF, while for CX3CR1-EGFP slices, AlexaFluor 594 carboxylic acid (ThermoFisher, Cat#A33082) was diluted to a final concentration of 200 μM. Images were acquired with a pixel size of 80 nm and pixel dwell time of 1.01 μs over a 125×125 μm^2^ field of view (FOV). For slices labeled with calcein, the pinhole was reduced to 0.5 Airy units (AU). For z-stacks, the step size was 0.5 μm, while for time series, the frame rate was 1 per 5 s for each 125×62.5 μm^2^ FOV. For slices labeled with AlexaFluor 594, the pinhole was increased to 1.0 AU. A lesion was induced by focusing laser illumination to a 2.5 μm^2^ region of interest for 1 s. Acquisition settings were reset to match baseline imaging parameters and subsequent images were acquired with a frame rate of 1 per 20 s for the 125 x125 μm^2^ FOV. The correction collar was set to a value that maximized image brightness and sharpness. Image acquisition was controlled by the Leica software.

### Light sheet microscopy

Muller organotypic slices were imaged using a custom-built lattice light sheet microscope (LLSM), which was a replica of the setup described previously ^14^. We used a dithered square lattice with an inner and outer NA of 0.44 and 0.55, respectively, giving a light sheet of constant thickness of approximately 0.6 μm over a length of 15 μm. Laser power incident on the illumination objective was controlled via an AOTF, and set to ∼60 μW for 10 Hz and ∼400 μW for 80 Hz acquisitions. The detection objective had an NA of 1.1 and a working distance of 2 mm. 3D volumes were acquired by a horizontal piezoelectric translator (step size 200 nm). The concentration of Calcein inside the imaging chamber was 20 μM. The camera frame rate was 10 Hz for **Fig. 2C** and **SI Video 5** and 80 Hz for **Fig. 2D** and **SI Video 6**. Image acquisition was controlled by home-made software written in LabVIEW (shared by HHMI via research license agreement).

### 2-photon microscopy (*in acute brain slices*)

Acute brain slices were imaged using a commercial 2-photon microscope (Prairie Technologies). For simultaneous two-color imaging of Alexa Fluor 488-Dextran (ECS) and YFP (tumor cells), the wavelength of the 2-photon laser (Ti:sapphire, Chameleon Ultra II, Coherent) was tuned to 920 nm, whereas for single-color imaging of Alexa Fluor 488, the wavelength was 850 nm. Images were acquired using a 40X water immersion objective with an NA of 1.0 (Plan-Apochromat, Zeiss). Laser power was between 10 to 25 mW in the focal plane. The fluorescence signal was spectrally divided into two channels by a dichroic mirror with a cut-off wavelength of 514 nm and collected by PMT detectors. Images were acquired with a pixel size of 144 nm over a 295×295 μm^2^ FOV and pixel dwell-times between 15-20 μs, corresponding to an acquisition time of 62 s per image. Image acquisition was controlled by the Prairie View software.

For local dye injections, we used patch pipettes. The borosilicate glass capillaries were pulled with a vertical puller (Narishige, PC-10, Japan) to a tip resistance between 3-5 MΩ and filled with an ACSF solution containing 500 μM Dextran-Alexa Fluor 488 dye (10,000 MW, anionic, fixable, Invitrogen). The pipette tip was placed some 50 to 80 μm below the slice surface to inject the dye below the damaged and cut open cells on the surface of the acute slice. 8 to 10 psi of pressure was applied via a Picospritzer III (Parker, Germany) for several minutes at a time, which yielded an even distribution of the dye within the FOV, which persisted long enough to acquire images with high contrast between ECS and cellular structures.

### 2-photon microscopy (*in vivo*)

Imaging was performed using a custom-built 2-photon microscope, based on a standard commercial upright research microscope (BX51WI, Olympus, Hamburg, Germany). 2-photon excitation was achieved using a femtosecond mode-locked fiber laser (Alcor 920, Spark lasers) delivering <100 fs pulses at a wavelength of 920 nm and a repetition rate of 80 MHz. Laser power was adjusted using a Pockels cell (302 RM, Conoptics) to around 20-30 mW power after the objective.

The wavefront of the laser beam was modulated by an SLM (Aberrior Instruments) to correct optical aberrations induced by the optics of the microscope and the sample. Appropriate lens combinations were used to conjugate the SLM on a telecentric scanner (Yanus IV, TILL Photonics), which projected both scan axes on the back focal plane of the objective lens (UPLanSAPO, 60X, silicone oil immersion, NA 1.3 Olympus) mounted on a z-focusing piezo-actuator (Pifoc 725.2CD, Physik Instrumente). The epi-fluorescence signal was descanned separated from incident beam using a long-pass dichroic mirror (580 DCXRUV, AHF) and detected by an avalanche photodiode (SPCM-AQRH-14-FC, Excelitas) with a bandpass filter (680SP-25, 520-50, Semrock) along the emission path. Signal detection and hardware control were performed with the Imspector scanning software (Abberior Instruments) via a data acquisition card (PCIe-6259, National Instruments).

### Assessing spatial resolution with fluorescent beads

Green fluorescent beads (diameter: 170 nm, Thermofisher) were dried on a coverslip and then mounted to a slide. 5 × 5 μm^2^ images were acquired on the respective microscopes under comparable imaging conditions. 5 μm thick z-stacks were acquired with 0.2 μm steps and the section in focus was selected to determine xy resolution. In-focus beads were imaged in xz format to determine axial resolution. Full width at the half maximum measurements were determined using the FWHM plug-in for FIJI.

### Image processing and data analysis

LISHI images were deconvolved in 3D with an open source Richardson Lucy (RL) algorithm (https://github.com/dmilkie/cudaDecon), using an experimentally measured PSF. The RL software is bundled into LLSpy, a python LLSM data processing and visualization toolbox (https://github.com/tlambert03/LLSpy). The deconvolution was run on a CUDA compatible GPU graphics card. A 3×3 median filter (noisy pixel removal) followed by background subtraction was applied before RL deconvolution with 20 iterations. Spurious horizontal and vertical stripes due to camera artifacts were removed with the ImageJ FFT filter. The deconvolved data was rotated to coverslip coordinates and resampled to obtain isotropic voxel dimensions (102^3^ nm^3^).

TUSHI images were visualized using ImageJ. Volume fraction of ECS was calculated from binarized images obtained either by wavelet analysis in SpineJ plugin^23^ (https://github.com/flevet/SpineJ) for ImageJ, either by custom made software in Python 3.0. OpenCV module (https://docs.opencv.org/4.x/) based on mean adaptive thresholding. Data was analyzed in MS Excel and statistical analysis was done in Graph Pad Prism 9. All data is represented as mean ± S.E.M.

## Notes

### Competing Interest Statement

The authors have declared no competing interest.

## References

1. Wang, Q. et al. The Allen Mouse Brain Common Coordinate Framework: A 3D Reference Atlas. Cell 181, 936–953 e920 (2020).

2. Abbott, L.F. et al. The Mind of a Mouse. Cell 182, 1372–1376 (2020).

3. Tonnesen, J., Inavalli, V. & Nagerl, U.V. Super-Resolution Imaging of the Extracellular Space in Living Brain Tissue. Cell 172, 1108–1121 e1115 (2018).

4. Hell, S.W. & Wichmann, J. Breaking the diffraction resolution limit by stimulated emission: stimulated-emission-depletion fluorescence microscopy. Opt Lett 19, 780–782 (1994).

5. Calovi, S., Soria, F.N. & Tonnesen, J. Super-resolution STED microscopy in live brain tissue. Neurobiol Dis 156, 105420 (2021).

6. Arizono, M., Inavalli, V., Bancelin, S., Fernandez-Monreal, M. & Nagerl, U.V. Super-resolution shadow imaging reveals local remodeling of astrocytic microstructures and brain extracellular space after osmotic challenge. Glia 69, 1605–1613 (2021).

7. Arizono, M. et al. Structural basis of astrocytic Ca(2+) signals at tripartite synapses. Nat Commun 11, 1906 (2020).

8. Velicky, P. et al. Saturated reconstruction of living brain tissue. biorxiv, 2022.2003.2016.484431 (2022).

9. Stoppini, L., Buchs, P.A. & Muller, D. A simple method for organotypic cultures of nervous tissue. J Neurosci Methods 37, 173–182 (1991).

10. Gahwiler, B.H. Organotypic monolayer cultures of nervous tissue. J Neurosci Methods 4, 329–342 (1981).

11. Hong, S., Dissing-Olesen, L. & Stevens, B. New insights on the role of microglia in synaptic pruning in health and disease. Curr Opin Neurobiol 36, 128–134 (2016).

12. Vainchtein, I.D. & Molofsky, A.V. Astrocytes and Microglia: In Sickness and in Health. Trends Neurosci 43, 144–154 (2020).

13. Keller, P.J. & Dodt, H.U. Light sheet microscopy of living or cleared specimens. Curr Opin Neurobiol 22, 138–143 (2012).

14. Chen, B.C. et al. Lattice light-sheet microscopy: imaging molecules to embryos at high spatiotemporal resolution. Science 346, 1257998 (2014).

15. Ducros, M. et al. in Neural Imaging and Sensing 2019 8 (SPIE, San Francisco, France; 2019).

16. Svoboda, K. & Yasuda, R. Principles of two-photon excitation microscopy and its applications to neuroscience. Neuron 50, 823–839 (2006).

17. Sakariassen, P.O. et al. Angiogenesis-independent tumor growth mediated by stem-like cancer cells. Proc Natl Acad Sci U S A 103, 16466–16471 (2006).

18. Daubon, T. et al. Deciphering the complex role of thrombospondin-1 in glioblastoma development. Nat Commun 10, 1146 (2019).

19. Kitamura, K., Judkewitz, B., Kano, M., Denk, W. & Hausser, M. Targeted patch-clamp recordings and single-cell electroporation of unlabeled neurons in vivo. Nat Methods 5, 61–67 (2008).

20. Nicholson, C. & Hrabetova, S. Brain Extracellular Space: The Final Frontier of Neuroscience. Biophys J 113, 2133–2142 (2017).

21. Miyai, M. et al. Current trends in mouse models of glioblastoma. J Neurooncol 135, 423–432 (2017).

22. Venkataramani, V. et al. Disconnecting multicellular networks in brain tumours. Nat Rev Cancer 22, 481–491 (2022).

23. Levet, F., Tonnesen, J., Nagerl, U.V. & Sibarita, J.B. SpineJ: A software tool for quantitative analysis of nanoscale spine morphology. Methods 174, 49–55 (2020).

